# Synchrotron-based DEI and DEI-CT systems to image chick pea seeds, plant anatomy and the associated physiology at 30 keV

**DOI:** 10.1101/2022.08.24.505216

**Authors:** D.V. Rao, G.E. Gigante, Z. Zhong, R. Cesareo, A. Brunetti, N. Schiavon, T. Akatsuka, T. Yuasa, T. Takeda

## Abstract

**Summary:**

- The study the effect of contrast on seeds, growth and the associated anatomy and physiology, with upgraded imaging systems. The use of phase information to explore new information at various stage of the growth.
- This work benefits, the use of Synchrotron-based DEI and DEI-CT systems to enhance the contrast in plant root architecture and contrast mechanisms, visibility of fine structures of root architecture growth and some aspects of physiology at acceptable level. These non-destructive, imaging systems available at the X-15A beamline, at NSLS, BNL, USA, are utilized.
- Noticed detailed anatomical and physiological observations, contrast mechanisms, with these upgraded systems, compared to other conventional techniques, equipped with tube source of X-rays. Examined the potential of these systems to quantify the plant roots in situ. The acquired images provided good contrast, anatomical structures and physiology of the plant root micro-architecture. We observed some of the complex plant traits, such as growth, development, root architecture and the associated physiology. The interior structure, root architecture, root morphology, growth of laterals and subsequent laterals can be visualized directly by synchrotron-based imaging techniques.
- Root architecture of the plant grown from seeds provides new information about the structure and enhancement of some desired property, for example, interior micro-structure of the root laterals and the subsequent laterals and the clear visibility of the leaves in detail. This way, it will be possible to differentiate the weakly and strongly attenuation of the signal traversing within the sample, clearly reflects the acceptable visibility in root laterals, subsequent laterals and the associated opaque matrix with enhanced contrast. The sample has a thin layer of hard structure outside and protein inside. Extinction properties of these samples will be characterized by Sy-DEI and Sy-DEI-CT. This way, we may be able to differentiate softly and weakly attenuation within the sample, to know more about the contrast mechanisms. The visibility, contrast and porosity, with finer details, can be noticed, with Sy-DEI-CT systems as distinguished from Sy-DEI. However, limited field of view, may limit the problems associated with Sy-DEI-CT.

## Motivation

Synchrotron-based techniques have been increasingly employed in various fields of physical, chemical, biological, geological, medical, material, and life sciences. Images from the samples demonstrate very weak absorption contrast, nevertheless producing significant phase shifts in the X-ray beam. Image processing derives structural features, which are then numerically quantified by image analysis. Contrast enhancement plays an important role in image processing; it enhances structural features that are hardly detectable to the human eye and allows automatic extraction of those features. The implementation of phase contrast based Sy-DEI and Sy-DEI-CT enables the investigation of the connectivity of the interior structure, which are invisible to the absorption contrast, while are readily observed in phase contrast. These imaging techniques are useful tools to examine the plant anatomy and physiology. Interaction between the light and plant, takes place, through reflected, absorbed or transmitted photons. Tomographic imaging with X-ray photons has been used in the non-destructive study of plant-soil interactions for a number of years. However, these techniques are very limited in use in the field of agriculture (Duliu, 1999; Gregory et al., 2003; Hamza et al., 2001; Heeraman et al., 1997; MacFall et al., 1994;2001; Moran., 2000; Perret., 1999;1999;2000;2003; Taylor., 1991; Timmerman., 2002; Tollner., 1994; Watanabe., 1992; Wulfsohn., 1999), food and measurement (Zwiggelaar., 1993., Rao et al., 2013). With CT system, a series of 2D images, from an object can be scanned, in a single axis of rotation and an assembly of 2D images, generates 3D image of an object. 3D volumetric data, from the growth of laterals and the subsequent laterals, provide the interior plant structures and root architecture. As regards to image analysis, ninety-two, different image analysis software tools are available, to study the plant biology (http://www.plant-image-analysis.org).

In recent years, improvements in image quality were carried out regularly on phantoms and biological samples with SY-DEI and SY-DEI-CT systems. Based on above and in continuation of our research, we have chosen Sy-DEI and Sy-DEI-CT systems to image seeds and their growth at different intervals of time. Limited studies have been reported using high energy X-ray imaging systems with synchrotron radiation, for imaging and quantifying plant root systems in situ (Young., 2007; Kao et al., 2007). A need therefore exists to evaluate these higher energy tomographic imaging systems with regards to resolution, noise and contrast. Since plant roots have higher water content, we hypothesize that, synchrotron-based systems are excellent tools for the observation and quantification of fine plant roots in situ. Plant roots serve as important organs for water and nutrient uptake, synthesis of growth regulators and storage of carbohydrates. In this investigation, we report on the potential use of the synchrotron-based imaging systems to study the nature of plant roots at a spatial resolution of about 9 µm or larger. On the other hand, non-destructive analysis is of great interest to a broad community of potential users in the field of agriculture and biological sciences (Lei et al.,2014).

Earlier researchers studied the plants by non-invasive image analysis (Leister et al., 1999), plant growth (Mühlich et al., 2008), mesh processing based technique for 3D plant analysis (Paproki et al., 2012), and maize roots (Grift et al., 2011). Further, various approaches, such as, 3D analysis of plant structure using X-ray CT (Stuppy et al., 2003), measuring root traits (Hargreaves et al., 2009), acquisition and analysis of CT scan data (Lontoc-Roy et al., 2006), non-destructive visualization and quantification of root using CT (Pierret et al., 2007) and visualization of undisturbed root architecture (Tracy et al., 2010) were examined extensively.

The spatial resolution of the present systems was 9 µm and has the potential of enabling entire root systems of seedlings to be reconstructed and their morphology. Further, in near future, to examine the growth rate, track development in the grown plant with anatomical mutations, imaging root growing in opaque matrix.

## The DEI system

The details of diffraction-enhanced imaging technology and the associated instrumentation have been presented previously (Rao et al. 2013) and we will high-light the X-ray optics scheme, with a brief description here. Fig.1. shows the schematic of the DEI system used for our experiments are available at the X-15A beamline, at NSLS, BNL, USA and for DEI-CT, the sample is placed on a Huber (Blake Industries, Scotch Plains, NJ) rotational stage. Highly collimated beam of X-rays are prepared by the Silicon [3, 3, 3] monochromator consisting of two perfect silicon crystals (Zhong, 2000). Once this beam passes through the subject, a third crystal (analyzer crystal) of the same reflection index diffracts the X-rays onto radiographic film (Kodak Professional Industrie 150, Industrex SR45), an image plate detector (Fuji HRV image plate, read out by a Fuji BAS2500 image plate reader at 50 micron pixel size) or a X-ray CCD detector (Photonic Science VHR camera, 9 micron pixel size).

**Fig 1.**
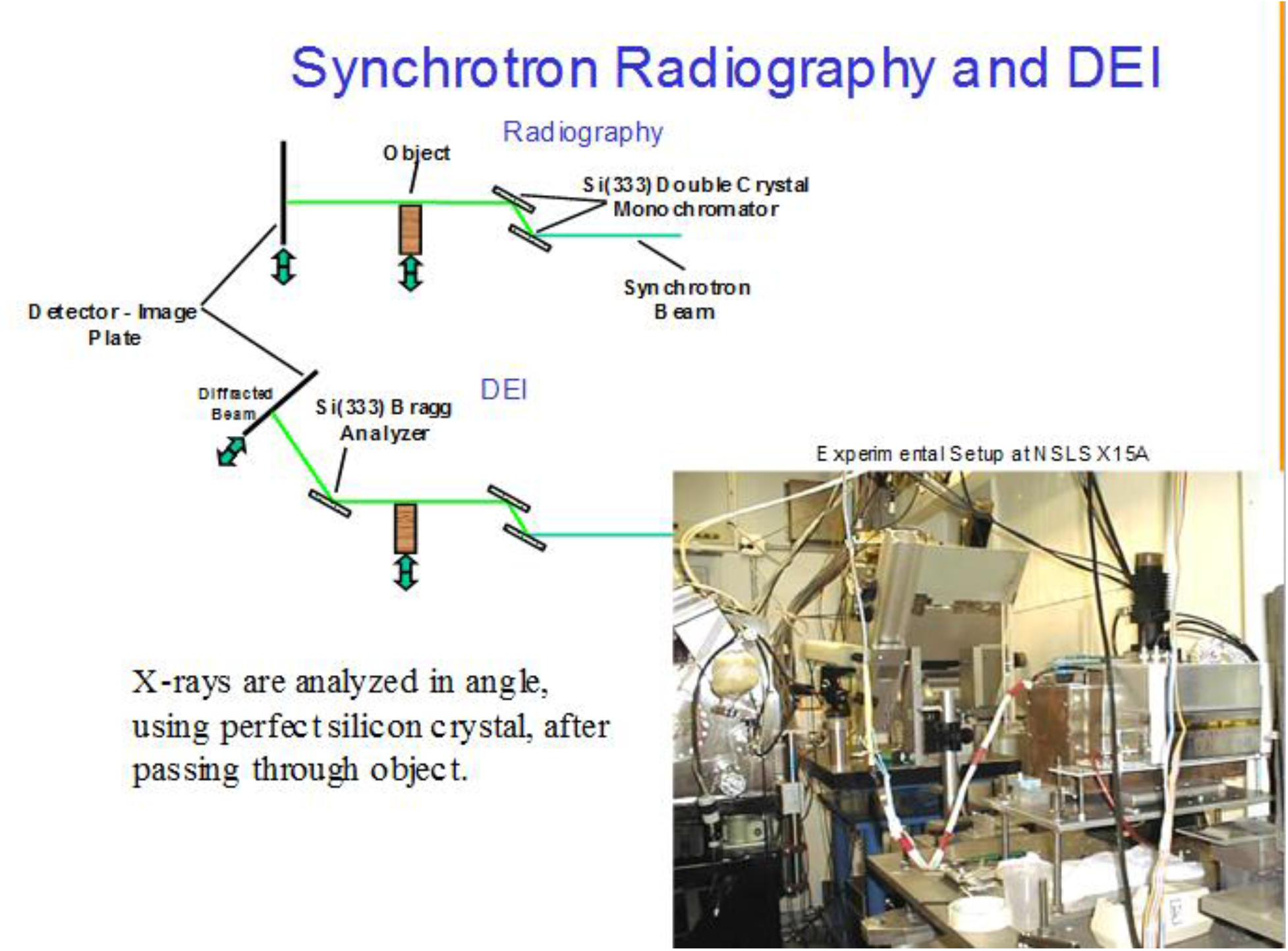
Experimental DEI and DEI-CT system.

## Samples

Chick pea seeds samples were washed for surface-sterilization in a weak solution of 10% sodium hypochlorite. Later the seeds were allowed to germinate on paper towels in distilled water for three days in a dark cabinet under optimum temperature conditions. Germination of the seeds was observed every day. The growth of the germination is imaged for every day and continued up to few days (Fig.2.).

**Fig 2.**
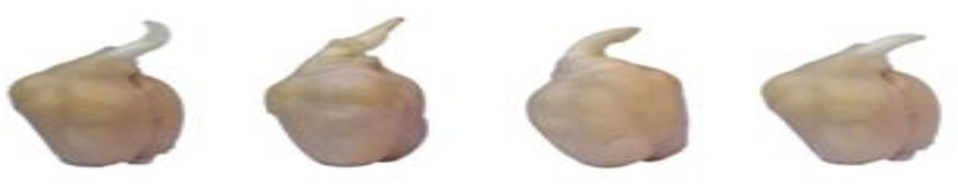
Growth of Chick pea seed after 24 hrs.

## Dose

DEI images are sharper than conventional X-ray images because higher energy X-rays (30 keV) are used and refracted X-rays are measured as data. An additional advantage of using high-energy X-rays is that the absorbed radiation dose is low. The amount of radiation absorption has been calculated to be < 1% (approximately 0.12 mGy per exposure). It will not affect germination or growth significantly.

## Data analysis

For the planar mode Sy-DEI imaging, the sample stage scanned the object vertically through the X-ray beam. These images were processed using the background subtraction tool in Image Pro Express 5.1 (Media Cybernetics, Bethesda, MD) and acquired 21images at different analyzer positions for each sample. Sy-DEI-CT scans were performed for the sample by fixing the analyzer crystal at a specific angle and rotating the sample stepwise through 360^0^ at 0.5^0^ or 18^0^ increment. The specific angle for the analyzer crystal was fixed at 11.62^0^. For each image in CT mode, 720 or 2,000 projections were acquired with a sample rotation step size of 0.5^0^ or 0.18^0^, and with an acquisition time of 1s per image. For tomographic reconstruction, a filtered back projection algorithm was used. Custom programs written in Interactive Data Language (IDL) were used to perform all of the image analysis, tomographic reconstructions and graphic visualization. The resolution of the detector is 9 µm but the reconstruction was performed with 2×2 rebinning to achieve a voxel size of 18 microns.

Beamline control and data collection were handled by a program called spec (Certified Scientific Software, Cambridge, MA, USA). The data was subsequently manipulated and images displayed using Igor Pro.5.04 (Wavemetrics, Lake Oswego, OR, USA). Each vertical column of image data was normalized, using a region of the image that did not contain any sample, to reduce differences in detector sensitivity. The average intensity and standard deviation of each pixel’s rocking curve determined and used in grey-scale image as a means of reducing the dimensionality of the data. The data were ordered into a three dimensional array, with the ‘x’ and ‘y’ dimensions corresponding to the spatial coordinates of the image and ‘z’ direction corresponding to the rocking curve angle, i.e. each of the 11, 21 or 31 points in the z direction represented by a different angle on the rocking curve (Chapman et al., 1997). The samples were imaged in both Sy-DEI planar and Sy-DEI-CT mode.

The seeds and developed seeds were placed in plastic container and fixed at the centre of the container. The laterals and the subsequent laterals are fixed in a hollow cylinder of 5mm inner diameter with a 2mm wall thickness, to avoid any artifacts, when the sample rotates. A silicon (333) analyzer crystal was placed after the sample stage and rotating using 1 µrad steps to capture only those photons striking it at the correct Bragg angle. Separated by 1 µrad steps, 11, 21, or 31 images were captured in addition to a background image which was made with the beam shutter closed. Photons reflected from the analyser crystal were captured using a Fuji HR-V image plate with a Fuji BAS-2500 reader, taking approximately 30 s to record each image.

## Results and discussion

Physical dimension of the chick pea seed and the formal growth is shown in fig.3. Fig.4. shows the growth rate of the seed, for twenty four hours, before acquisition of the image with Sy-DEI and Sy-DEI-CT. Fig.5. shows the, acquisition of the image in planar mode with Sy-DEI and the associated histograms, reflect the density of the plant growth at various parts. The mesh structure of the leaf at the central position is less dense as distinguished from the extreme ends. The embedded root architecture can be observed in the image, since the sample refracts X-rays to a greater extent than the surrounding matrix. Darker pixels in the attenuation image indicate regions where greater refraction or scatter occurred. Histogram’s, with roots hidden at left side, reflects on the density on the density of the growth rate of the laterals and the subsequent laterls.

**Fig 3.**
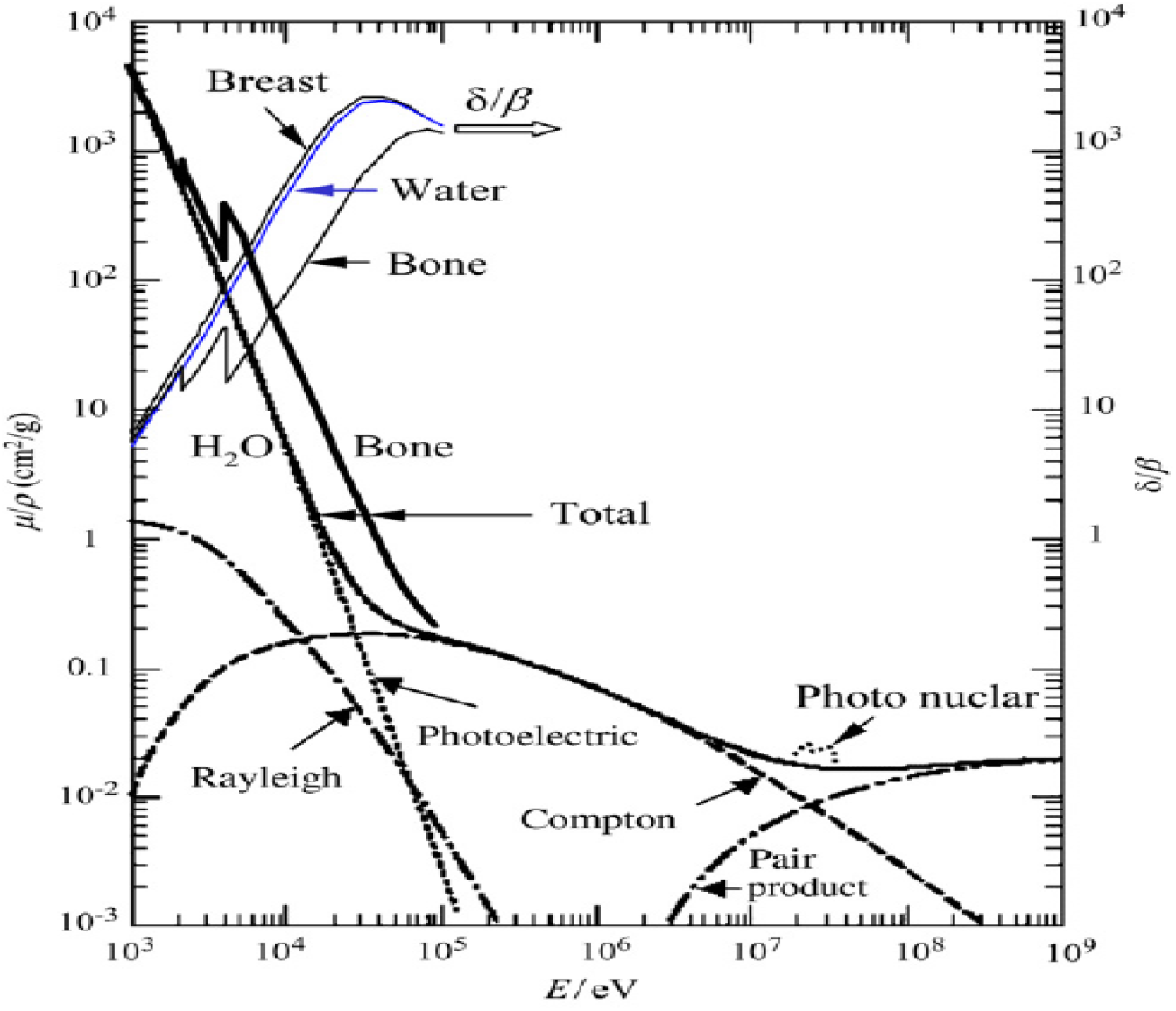
The dependence of phase to attenuation contrast ratio for few soft biological samples.

Sy-DEI and Sy-DEI-CT, provided clearer, less noisy images, than those obtained using conventional X-ray imaging. The seed growth is denser than parts of the seed architecture and thus caused significant x-ray refraction. Further analysis of the data could provide useful information about the amount of the refraction and scatter resulting at each pixel, which could then be interpreted as relative density and concentration of ordered molecules, respectively.

The physiology and the associated morphology of these soft materials provide the crucial tool for determining the structure and enhancement of some desired property, for example, soft external layer and interior structure. It will be possible to differentiate the weakly and strongly attenuating materials. The has soft external layer, having some having hard inside for the growth of the plant and the associated physiology. This way, we may be able to differentiate softly and weakly attenuating materials, to know more about the contrast mechanisms. A wide range of samples are studied in medicine, bio-medical, biological and material science, which can produce significant phase shifts of the X-ray beam.

The image features are shown with a scale bar in fig.6. We have chosen the beam profile, projection area, are at the center of the sample. The intensity profile is symmetric, towards the low-angle, high-angle and at the peak position. Fig.7. shows the intensity variation at positions of the image. Fig.8.(a) to (d) shows, Sy-DEI-CT refraction images of the sample, towards the low angle, at the peak position and high-angle side, profile and the rendered image of the sample at 30 keV. The images shows the effectiveness of the tomography mode for better visualization. It is interesting to note that, in the energy region from 15 to 30 keV, the phase shift term ‘δ’ is of the order of 10^−7^ and the absorption term ‘β’ is of the order of 10^−10^, reflecting to reveal the phase effects even if the absorption contrast is low. The contrast has considerable visibility in the tomographic mode. The reconstructed image of the root architecture, with a nature of porosity of the root interior structure, with a wide hallows nature. The problem with tomography is that a large number of images must be captured, resulting in greater radiation exposure. Radiation exposure with two-dimensional DEI is low. The effects of radiation damage can be reduced by reducing exposure times and frequencies or by reducing the number of data points included in the rocking curve.

Comparison of the reconstructed root image ((Fig.8 (a) to (d)) with (Fig. 4), it is obvious that the number of laterals and the subsequent laterals have same dimensions of roots and the associated morphology of the plant growth. Furthermore, fine roots with diameters less than 0.5mm are detected and all artefacts from the background and sample container are removed. This is of great importance for a consecutive analysis mapping the laterals and the subsequent laterals of the root architecture system at 9µm resolution. Fig’s. 8(a) to (c) show the processed CT images. These images were obtained, after directly thresholding the filtered data.. The same threshold level is applied to the volume rendering (fig.8(e)).

**Fig 4.**
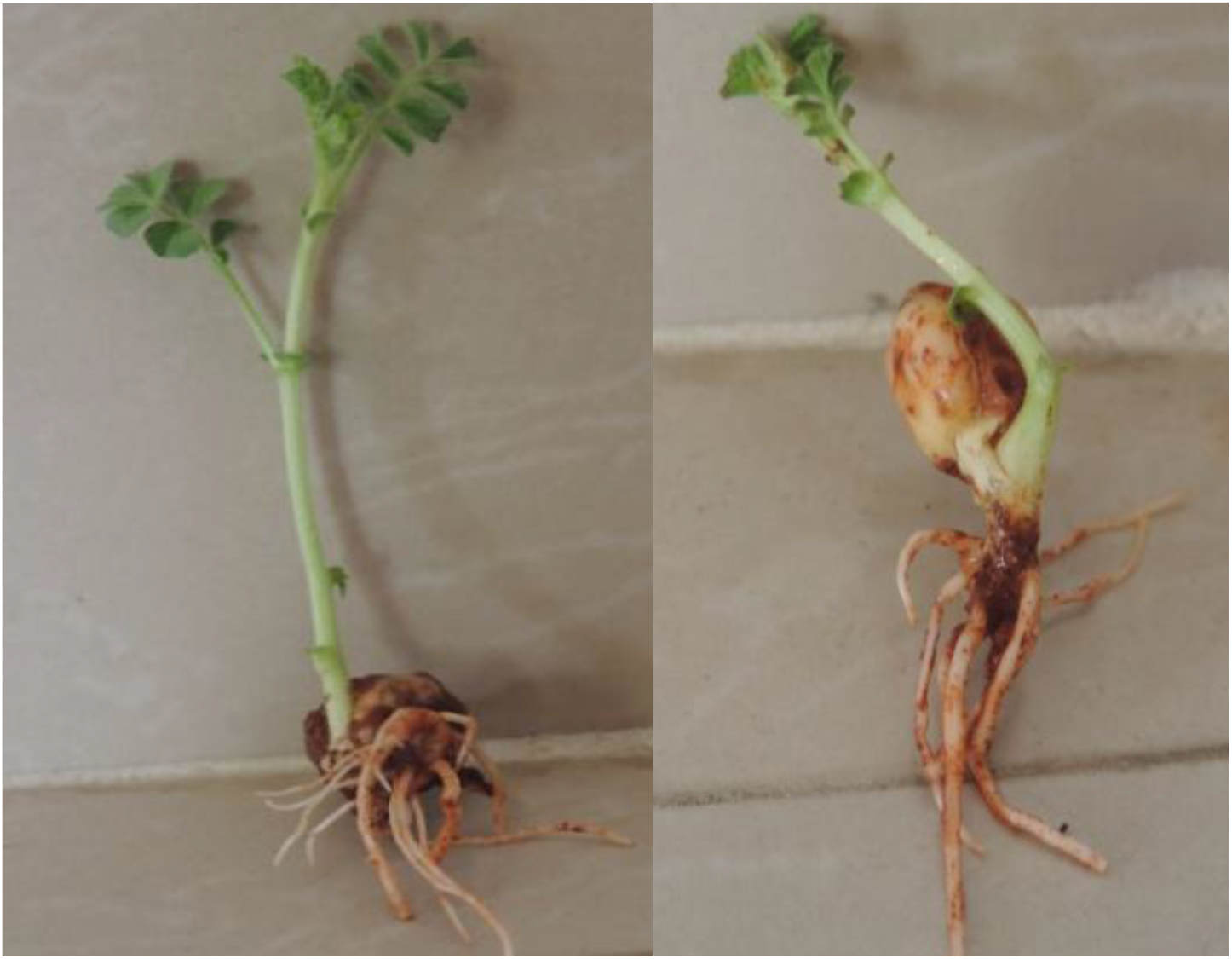
The root laterals, subsequent laterals, and their lengths, growth of the seed before the acquisition of the images and the roots used for CT image reconstruction. The length of the root varies from 2 to 5cm and the diameter varies approximately from 2 to 1.5 mm.

**Fig 5.**
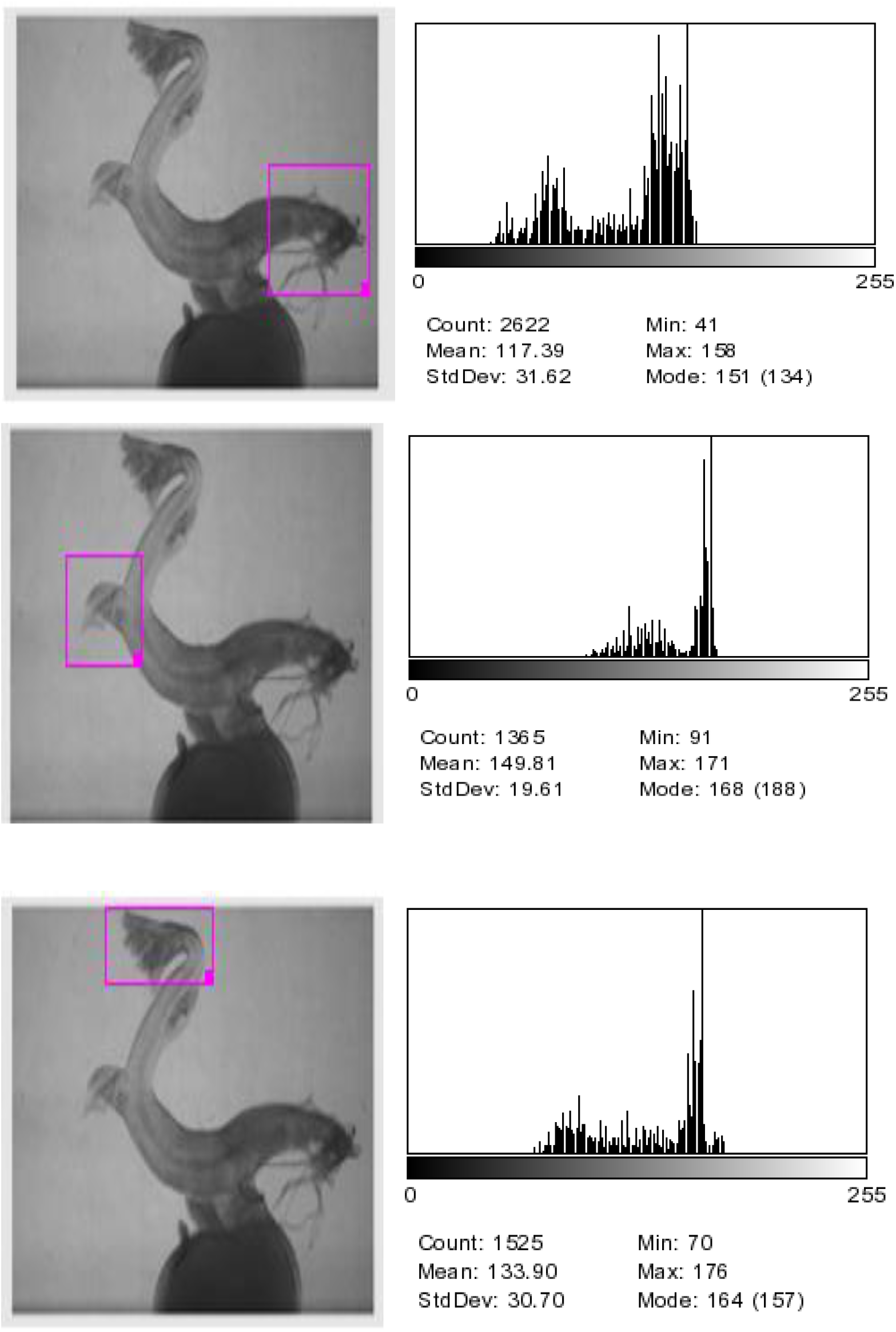
DEI images of the plant with leafs, growth, root architecture and the associated histograms, in the marked areas of the leafs, in planar mode.

**Fig 6.**
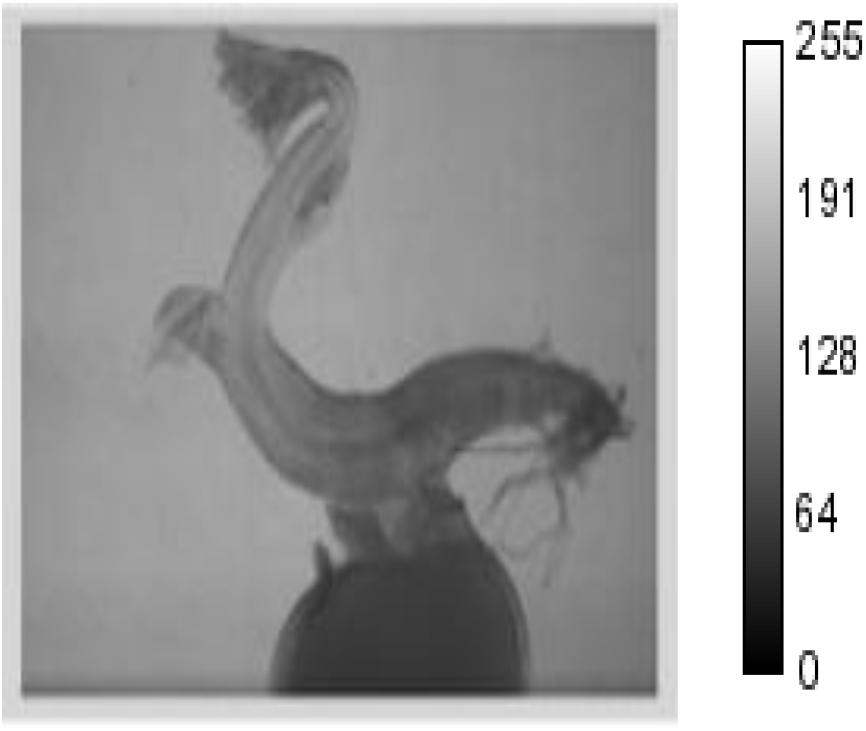
DEI with scale bar.

**Fig 7.**
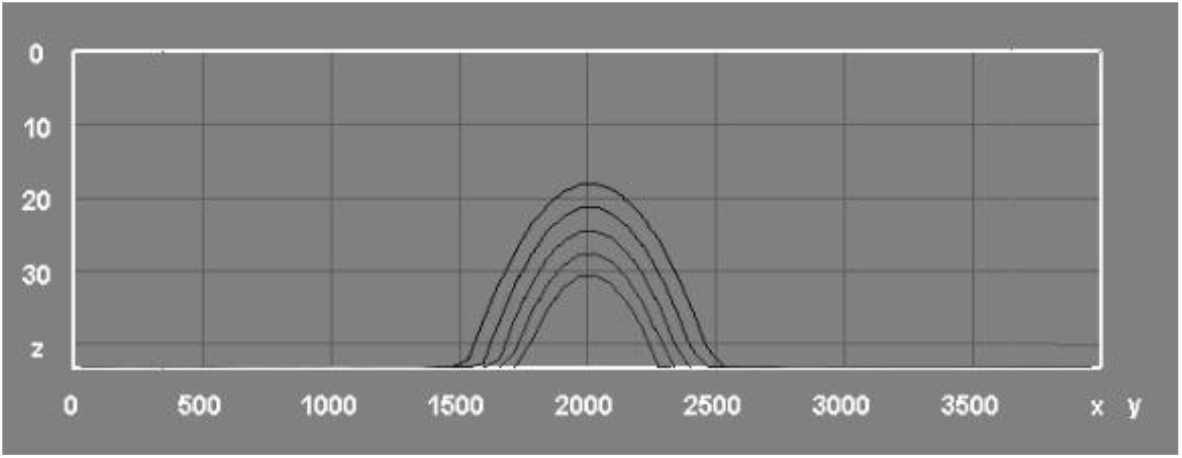
Interior variation of the intensity.

**Fig 8.**
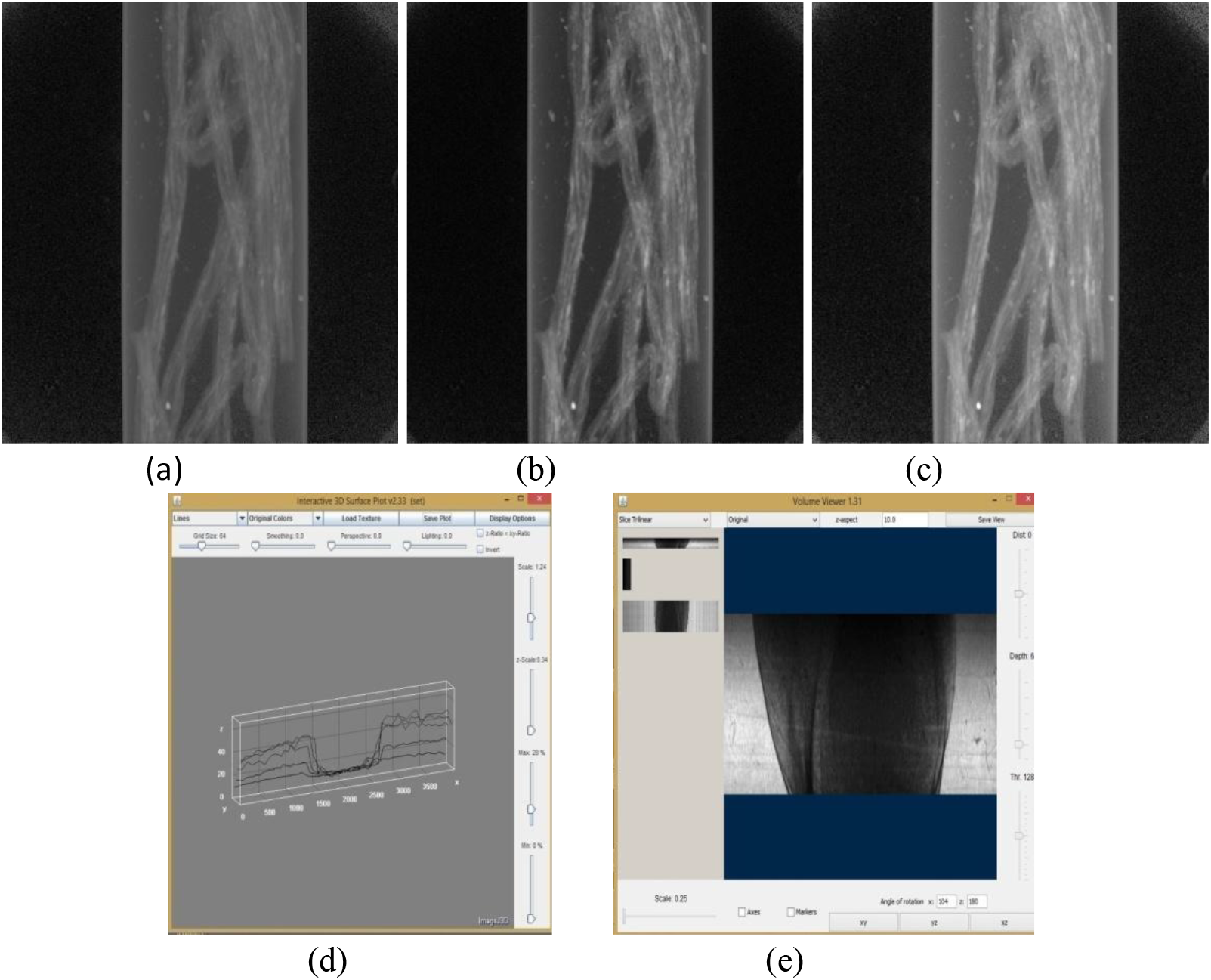
Internal root architecture structure of the sample in CT mode. Filtered CT images (a) Image at low-angle side (b) Image at the center (c) Image at high-angle side of the root (d) Intensity profile (e) Rendered image of the sample.

The systems and the associated software, allowed us, successfully, to reconstruct the fine root architecture of the laterals and subsequent laterals in Sy-DEI-CT. For the present image analysis and the scanning resolution of the Sy-DEI-CT system, it will be possible to observe and detect processes of the fine root architecture of laterals and the subsequent laterals, which are essential for plant growth. With the development of new digital technology, algorithms, it will easy to visualize the interior root architecture of the laterals and the subsequent laterals, with more visibility. At the end, more than 90% of the root volume of the examined sample and the root architecture, can be achieved at 9µm resolution.

## ACKNOWLEDGEMENTS

One of the author’s (DVR) undertook part of this work with a support from, Department of Science Based Applications to Engineering, Universita di Roma “La Sapienza”, Via Scarpa 10, 00161, Roma, Italy, Istituto di Matematica e Fisica, Universita di Sassari, Italy, Department of Bio-Systems Engineering, Yamagata University, Yonezawa, Japan and in the form of collaboration form the beamline scientist (Zhong Zhong), NSLS, BNL, USA “Use of the National Synchrotron Light Source, Brookhaven National Laboratory, was supported by the U.S. Department of Energy, Office of Science, Office of Basic Energy Sciences, under Contract No. DE-AC02-98CH10886”.

